# Dissociable roles of neural pattern reactivation and transformation during recognition of words read aloud and silently: An MVPA study of the production effect

**DOI:** 10.1101/2024.02.20.581164

**Authors:** Lyam M. Bailey, Heath E. Matheson, Jonathon M. Fawcett, Glen E. Bodner, Aaron J. Newman

## Abstract

Recent work surrounding the neural correlates of episodic memory retrieval has focussed on the decodability of neural activation patterns elicited by unique stimuli. Research in this area has revealed two distinct phenomena: (i) neural pattern reactivation, which describes the fidelity of activation patterns between encoding and retrieval; (ii) neural pattern transformation, which describes systematic changes to these patterns. This study used fMRI to investigate the roles of these two processes in the context of the production effect, which is a robust episodic memory advantage for words read aloud compared to words read silently. Twenty-five participants read words either aloud or silently, and later performed old-new recognition judgements on all previously seen words. We applied multivariate analysis to compare measures of reactivation and transformation between the two conditions. We found that, compared with silent words, successful recognition of aloud words was associated with reactivation in the left insula and transformation in the left precuneus. By contrast, recognising silent words (compared to aloud) was associated with relatively more extensive reactivation, predominantly in left ventral temporal and prefrontal areas. We suggest that recognition of aloud words might depend on retrieval and metacognitive evaluation of speech-related information that was elicited during the initial encoding experience, while recognition of silent words is more dependent on reinstatement of visual-orthographic information. Overall, our results demonstrate that different encoding conditions may give rise to dissociable neural mechanisms supporting single word recognition.

## 1. Dissociable roles of neural pattern reactivation and transformation during recognition of words read aloud and silently: An MVPA study of the production effect

Over the past two decades, developments in multivariate analyses of functional neuroimaging data have enabled researchers to decode information distinguishing individual stimuli from one another, based on the unique patterns of neural activation that they elicit (Peelen and Downing, 2022). This approach has been of great value to research examining neural correlates of episodic memory retrieval. A major finding in this area is that the fidelity of activation patterns (within a particular brain area) to a unique study item between encoding and retrieval reliably predicts retrieval success for that item (Davis et al., 2014; Ritchey et al., 2013; Kuhl and Chun, 2014; Wing et al., 2015). This *reactivation* often occurs in high-level visual processing centres such as ventral temporal and lateral occipital cortices, reflecting perceptual processing demands that are present during encoding (e.g., studying word-picture pairs) but absent during retrieval (e.g., when viewing the word alone; Jafarpour et al., 2014; Danker et al., 2017; Favila et al., 2018). Neural reactivation might therefore be attributed to faithful mental reinstatement of the episode, grounded in the perceptual experiences that the episode entailed (e.g., Meyer and Damasio, 2009).

In contrast to reactivation, neural *transformation* describes systematic alteration of activation patterns elicited by the same stimulus. To our knowledge, transformation has been captured by prior literature in three main ways. First, information that is decodable in perceptual cortices (e.g., visual areas) during encoding may be reinstated in frontoparietal regions during retrieval (Xiao et al., 2017, 2020; Favila et al., 2018). Transformation has also been captured as a change in activity patterns within a discrete region; one may measure the similarity of responses between two encoding episodes, and then again between encoding and retrieval. A decrease in similarity between encoding and retrieval (relative to encoding-encoding)—but without a loss of item-specific decodability at retrieval—reflects transformation within that region (Chen et al., 2017; Xiao et al., 2017). Importantly, this second definition of transformation does not constitute a mere weakening of reactivation effects or an increase in noise, because item-specific information can still be decoded from the transformed neural patterns (e.g., Chen et al., 2017; but see Xue, 2022 for review and discussion). This kind of transformation may therefore be conceptualised as a change in the representational geometry (Kriegeskorte and Kievit, 2013) of stimulus-elicited activation patterns within a given region. Finally, transformation has also been captured as a change in the representational format of encoded information (i.e., the decodability of stimulus properties in activation patterns; Kriegeskorte et al., 2008). For example, some studies have characterised transformation as a shift from decodability of perceptual information to semantic information (Linde-Domingo et al., 2019; Liu et al., 2021). Broadly, transformation (by any of the above definitions) is thought to reflect reorganisation of encoded information, either as part of the memory consolidation process or to meet specific retrieval goals (Favila et al., 2020; Xue, 2022). This perspective is consistent with the general view that remembering is more of a reconstructive than reproductive process (Schacter et al., 1998).

Reactivation and transformation may be relevant to the *production effect*, which is a robust recognition advantage for words read aloud compared to words read silently (MacLeod et al., 2010). Much work in this area emphasises the role of sensorimotor experiences conferred by verbalisation: movements of the tongue and jaw, vibration of the larynx, and auditory feedback from one’s own voice. A popular view is that these distinctive experiences are appended to the episodic representation for having seen a word, and subsequently may be leveraged to facilitate retrieval. For example, one account holds that, during retrieval, participants may recollect (i.e., mentally replay) the productive act; this enables them to reject or endorse the presented word (e.g., on an old/new judgement task) using a so-called distinctiveness heuristic: *I remember reading this word aloud, therefore I probably studied it in this experiment* (Taikh and Bodner, 2016; MacLeod and Bodner, 2017). In a similar vein, Wakeham-Lewis et al. (2022) recently proposed that when participants are presented with a word during a recognition task, they may mentally simulate reading that word aloud; the simulated experience may then be compared with the encoded episode to endorse or reject the presented word. Importantly, both of these accounts depend on access to encoded sensorimotor information that is elicited when reading aloud. At a neural level, both perspectives seem compatible with a reactivation-based perspective whereby retrieval entails recapitulation of perceptually-grounded experiences. That is, if distinctive sensorimotor experiences are mentally replayed or simulated (and then compared) at retrieval, retrieval ought to entail reactivation of neural patterns elicited during speech production. Remembering distinctive speech experiences may also entail neural transformation. Using a distinctiveness heuristic (MacLeod and Bodner, 2017), or comparing mentally simulated actions to encoded experiences (Wakeham-Lewis et al., 2022), may entail evaluation an/or reorganisation of encoded sensorimotor information—this would be consistent with work showing that the decodability of neural responses to the same stimulus can vary according to one’s task (Nastase et al., 2017; Favila et al., 2018; Wang et al., 2018). Stated differently: speech-elicited information *for encoding* might manifest differently than *for retrieval*.

The goal of this study was to characterise how reactivation and transformation contribute to the production effect, using a whole-brain searchlight method^1^. Given the novelty of this work (to our knowledge, no prior work has investigated either process in the context of speech production), we were concerned that basing our analyses on a-priori regions of interest (ROIs) might blind us to potential areas of reactivation or transformation. As such, using a searchlight enabled fully data-driven identification of regions exhibiting interesting effects.

Participants read words aloud or silently (encoding phase), and later performed a recognition test (recognition phase), during fMRI scanning. We first computed behaviourally-relevant measures of reactivation and transformation using a whole-brain searchlight approach, for each individual subject. We quantified transformation as a systematic decrease in within-region pattern similarity (i.e., the second method of capturing transformation described earlier). Importantly, this measure was constrained by a between-subjects transformation analysis (similar to Chen et al., 2017) which ensured that results were not driven by an increase in noise between encoding and recognition. Both measures were behaviourally relevant in that they quantified reactivation or transformation for subsequently remembered words relative to forgotten words; hence, they revealed areas in which either process was associated with successful recognition. We then compared, at the group level, reactivation and transformation between the aloud and silent reading conditions.

We predicted that, relative to words that were read silently, words that were read aloud would elicit reactivation in primary and associative sensorimotor and auditory cortices linked to speech production (Dietz et al., 2005; Bailey et al., 2021; Qu et al., 2022). Transformation effects may be present in these same speech-relevant areas, or alternatively in frontoparietal areas implicated by prior work on transformation between encoding and retrieval (Chen et al., 2017; Xiao et al., 2017, 2020; Favila et al., 2018). Although our hypotheses were mainly focussed on processes that were increased for aloud words (revealed by aloud > silent contrasts), it was also possible that reactivation or transformation might be increased during recognition of silent words (silent > aloud). Therefore, our analyses entailed bidirectional between-condition contrasts of reactivation and transformation.

## 2. Methods

### 2.1. Subjects

Thirty participants took part in this experiment, aged 18-40 (*M* = 21.43, *SD* = 4.57), 21 female, three left-handed. Participants were recruited through on-campus advertising at Dalhousie University, and received $30 CAD reimbursement and a digital image of their brain. Prior to taking part in the study, all potential participants were screened to ensure normal or corrected-to-normal vision, proficiency in English, no history of neurological illness or trauma, and no contraindications to MRI scanning. Handedness information was obtained using the Edinburgh Handedness Inventory (Oldfield, 1971). The experiment took place at the IWK Health Centre in Halifax, Canada, and all procedures were approved by the IWK Research Ethics Board. Participants provided informed consent according to the Declaration of Helsinki.

Of the 30 participants recruited, five were excluded from data analysis. One participant could not complete the experiment due technical problems; another disclosed that they did not fit inclusion criteria (neurological normality) after having completed the study. Two participants reported problems or errors completing the task as instructed: one reported that they silently mouthed words in the aloud condition instead of vocalising; the other had difficulty reading the presented words in the scanner. Finally, one participant made zero incorrect responses in at least one condition during the recognition phase, making them ineligible for our measures of reinstatement and transformation (which depended on contrasts between correct and incorrect responses). Therefore, data from 25 participants (17 female, 2 left-handed, *M* age = 21.40, *SD* = 4.73) are reported here.

### 2.2. Stimuli and Apparatus

Participants viewed all stimuli while lying supine in the MRI scanner; stimuli were presented using the VisualSystem *HD* stimulus presentation system (Nordic Neuro Lab, Bergen, Norway). During the experiment, participants made button-press responses using MR-compatible ResponseGrip handles (Nordic Neuro Lab), one in each hand. Stimuli were presented using PsychoPy 2020.2.1 (Peirce et al., 2019); these appeared on an LCD screen positioned behind the scanner and viewed by participants via an angled mirror fixed to the MR head coil. Stimuli for the encoding and recognition tasks were 90 nouns selected from the list used in Bailey et al. (2021)^2^, shown in Table 1. Words were 6 to 10 characters in length, each with a frequency greater than 30 per million (Thorndike and Lorge, 1944). Stimuli for the active baseline task (see below) were printed numbers 1-9. All stimuli (words and numbers) were presented at the centre of the screen, in white lowercase Arial font against a dark grey background (RGB: 128, 128, 128). During the encoding phase, participants were asked to respond to each presented word by either reading it aloud or reading it silently. Response cues were greyscale icons either depicting a mouth (read aloud) or an eye (read silently), presented at the centre of the screen.

**Table 1.** Stimulus words used in the current study.

|  |  |  |  |  |  |  |  |  |
| --- | --- | --- | --- | --- | --- | --- | --- | --- |
| address | branch | distance | glass | justice | merchant | plate | shadow | treasure |
| afternoon | building | election | gravity | kettle | minute | pocket | shoulder | uncle |
| amount | capital | engine | guardian | kitchen | neighbour | porch | station | uniform |
| arrow | captain | entrance | harbour | knock | nephew | quarrel | steam | vacation |
| attention | castle | evening | history | ladder | office | quarter | stream | victory |
| attitude | clothes | factory | holiday | laugh | orchard | queen | teacher | village |
| avenue | daughter | fashion | invention | leather | package | resort | theatre | wagon |
| basket | debate | foundation | invitation | lesson | pants | reward | thread | wheel |
| battery | dinner | friend | island | market | partner | river | traffic | whisper |
| border | direction | furniture | judge | meadow | peace | school | travel | winter |

### 2.3. Procedure

Participants first provided informed consent and completed MRI safety screening and the Edinburgh Handedness Inventory. Participants were informed that they would be taking part in a study on memory, and that they would be studying a list of words that they would later be tested on. Before entering the scanner, participants completed a shortened practice version of the main experiment, using a different stimulus set, on a laptop computer to familiarise them with the experimental tasks. After this practice task, a member of the research team confirmed (verbally) that participants understood the task instructions. Following this, participants were positioned in the MRI scanner. A brief scout scan determined head position. Participants then completed three functional scans followed by a structural scan. The first two functional scans (approximately 13 minutes each) comprised the *encoding phase*; the third (approximately 10 minutes) comprised the *recognition phase*.

At the start of each participant’s scanning session, the 90 stimulus words (Table 1) were randomly allocated to one of three conditions: read aloud, read silently, or foil (30 words per condition). Words in the first two conditions were presented throughout the encoding phase, while all three conditions were present in the recognition phase. Stimulus-condition mappings were randomised between participants.

#### 2.3.1. Encoding Phase

Each functional run of the encoding phase comprised 60 trials, and each word (from the read aloud and read silently conditions) was presented once per run (this meant that each word was seen twice during the study phase). Word presentation order was randomised for each run. Each trial began with a fixation cross (“+”) presented for 500 ms, followed by a 1000 ms cue instructing participants to read the upcoming word either aloud or silently. The stimulus word for that trial was presented for 2500 ms, during which time participants were required to read the word as instructed. Following this, participants completed an active baseline task for 8000 ms. During the baseline task, a randomly generated number 1-9 appeared onscreen, and participants were instructed to indicate with a button press whether the number was odd or even. Response mappings (left-hand button press for odd, right-hand for even) were presented in the top corners of the screen and were consistent across all participants. Each number was presented for 2000 ms, after which it was replaced with a different (randomly generated) number; this procedure was repeated until the end of the trial. Thus, the active baseline task for each trial comprised odd-even decisions on four numbers, each presented for 2000 ms. This procedure resulted in a trial SOA of 12 seconds. A schematic of encoding phase trial structure is shown in the top panel of Figure 1.

**Figure 1.**
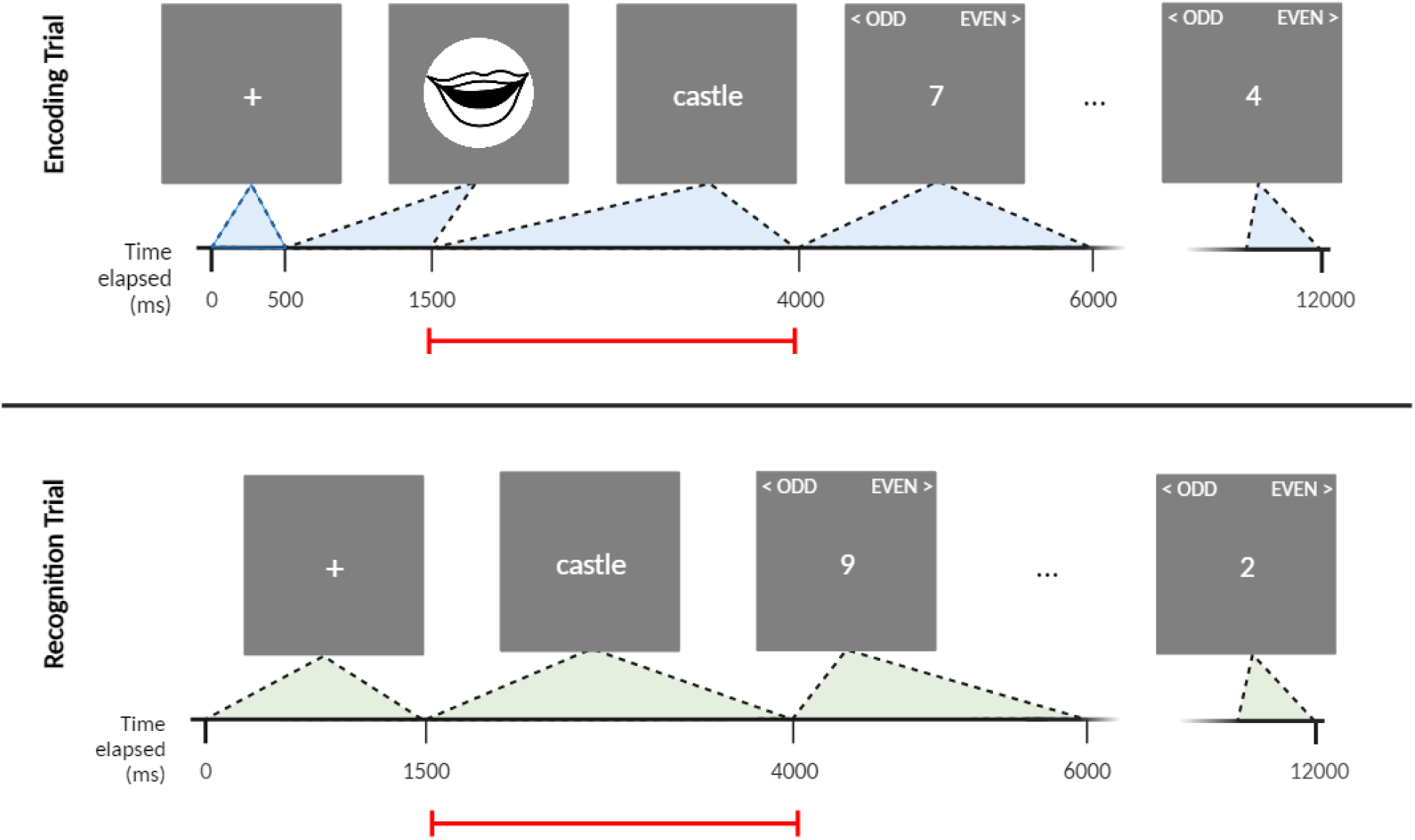
Schematic of an example trial from the encoding phase (top panel) and recognition phase (bottom panel). The red line in each panel indicates the temporal window modelled by the time series regressor for each trial (i.e., the temporal window from which activation patterns were estimated).

#### 2.3.2. Recognition Phase

The recognition phase comprised a single run with 90 trials, in which all stimulus words were presented in random order. Sixty of these words were previously seen by participants in the encoding phase (30 aloud; 30 silent); the remaining 30 were unseen foils. Each trial began with a fixation cross for 1500 ms, followed by a stimulus word for 2500 ms. During this 2500 ms period, participants were instructed to indicate with a button press whether the presented word was “old” or “new”. Response mappings (left hand for old, right for new) were consistent across participants, and cues were presented in the top corners of the screen as reminders. The study word remained onscreen for the full 2500 ms regardless of when participants made a response. Following this, participants completed an identical active baseline task to that of the encoding trials before the start of the next trial. This procedure ensured a trial SOA of 12 seconds, as in the encoding phase. A schematic of recognition phase trial structure is shown in the lower panel of Figure 1.

#### 2.3.3. MRI Data Acquisition

MRI data was acquired on a 1.5 Tesla GE MRI system (GE Medical Systems, Waukesha, WI) equipped with a 19 channel head coil. Each participant completed three functional scans (described above) followed by a structural scan. Functional scans used a gradient echo-planar pulse sequence, TR = 1800 ms, TE = 23 ms, FOV = 240 mm, flip angle = 90°. Images were obtained in 34 axial slices^3^ (no gap, sequential acquisition) of thickness = 3.75 mm, matrix = 64 x 64, resulting in an in-plane voxel resolution of 3.75 x 3.75 mm. The FOV included full cortical coverage and partial cerebellar coverage. We collected 400 functional volumes for each encoding run; 600 for the recognition run. Five additional volumes were acquired at the start of each run, but were discarded following acquisition. For the structural scan, a T1-weighted anatomical image was obtained using a magnetization-prepared rapid acquisition gradient echo (MPRAGE) sequence, TI = 1134 ms, flip angle = 8°, NEX = 2, FOV = 224 mm, matrix = 224 x 224, resulting in an in-plane voxel resolution of 1 x 1 mm.

### 2.4. fMRI Data Processing

fMRI data were preprocessed using functions from FEAT (fMRI Expert Analysis Tool) Version 6.00, part of FSL (FMRIB’s Software Library, www.fmrib.ox.ac.uk/fsl). FSL functions were implemented with custom bash shell scripting, in parallel for multiple subjects/runs using GNU *parallel* (Tange, 2011) to improve efficiency. Preprocessing steps included non-brain removal with BET (Smith, 2002), grand-mean intensity normalisation of the entire 4D dataset by a single multiplicative factor, high pass temporal filtering (Gaussian-weighted least-squares straight line fitting, with sigma = 50.0 s), spatial smoothing (FWHM Gaussian kernel, 6 mm), and motion correction using MCFLIRT. Our a priori threshold for excessive head motion between contiguous time points was 2 mm; no participant exceeded this threshold on any run.

We extracted trial-level activation patterns using the least-squares-all (LSA) method. Using FEAT, we modelled each of the three functional runs using a GLM with one regressor per trial. Each regressor comprised a time series modelling the period during which the word for that trial was presented (duration: 2500 ms), convolved with a gamma function (lag: 6 sec, sigma: 3 s) as a model of the haemodynamic response function (HRF). Each GLM also included parameters from motion correction as regressors of no interest. For each model, we defined contrasts of interest whereby each contrast comprised one regressor (i.e., trial) in the model. As a result, each participant was associated with 210 whole-brain functional maps (contrasts of parameter estimates; COPEs) in native functional space, with each map corresponding to a unique trial (60 in each encoding run; 90 in the recognition run). We next spatially transformed these functional maps to the MNI152 template. First, each participant’s example functional volume (that is, a functional image acquired in the middle of each run) was rigidly aligned to their high-resolution structural T1 image. Example functional volumes were taken from whichever run the to-be-transformed functional map belonged to. Each structural image was also aligned to MNI152 space with an affine transform. We then combined these EPI-to-structural and structural-to-MNI152 transforms to generate a matrix for transforming data from native functional space to MNI152, for each subject and run. We applied these matrices to the functional maps derived from the GLMs, such that each participant was associated with 210 functional maps in MNI152 space, re-sliced to 2 mm isotropic resolution. These transformation steps were performed using FSL’s FLIRT function (Jenkinson and Smith, 2001; Jenkinson et al., 2002). For convenience, these preprocessing stages were applied to words in all conditions, however we discarded output for foil words (from the recognition phase) from subsequent fMRI analyses.

### 2.5. Subject-Level Analyses

We computed behaviourally-relevant measures of reactivation and transformation (described in the following sections) and implemented each using a whole-brain searchlight (spherical searchlight area, radius = 3 voxels, average volume = 115.4 voxels). These measures were behaviourally-relevant in that they quantified the difference in reactivation or transformation between subsequently remembered and forgotten items. Both measures considered correlations between activity patterns in the same local region (i.e., same searchlight sphere) across encoding and recognition. Searchlights for each measure were implemented independently for each participant and condition (aloud or silent). The searchlights and all other procedures described in this section were carried out in the MATLAB environment using functions from the CoSMoMVPA toolbox (Oosterhof et al., 2016) and custom scripting.

#### 2.5.1. Reactivation

Reactivation analyses were based on the approach used by Zeithamova et al. (2017). These authors devised a measure for what they described as a ‘subsequent memory effect’; this measure was intended to capture larger same-item pattern similarity for subsequently remembered stimuli relative to subsequently forgotten stimuli, as observed in previous literature (see Introduction). For convenience, we will refer to this measure in our study as a *reactivation index*, recognising that it still captures the relationship between pattern similarity and subsequent memory. First, we sorted activation patterns (contrasts of parameter estimates from the GLMs described in 2.5) according to the run (Encoding 1, Encoding 2, or Recognition) and stimulus to which they corresponded. Next, in each searchlight sphere, we computed pairwise correlations between patterns elicited by the same item across both encoding runs and the recognition run. That is, for each item, we computed *r*(Encoding 1, Recognition) and *r*(Encoding 2, Recognition); correlation values derived from both encoding runs were included in the reactivation index calculation (described below). All correlation values were sorted according to whether the eliciting stimulus was subsequently remembered (correctly responded “old” during the recognition task) or forgotten (incorrectly responded “new”) by that participant. Following this, we computed a reactivation index as the standardised mean difference (Cohen’s D) between remembered same-item correlations and forgotten same-item correlations. This calculation is illustrated in Figure 2. This procedure yielded a whole-brain map of reactivation indices, and was implemented independently for each participant and condition (aloud or silent).

**Figure 2.**
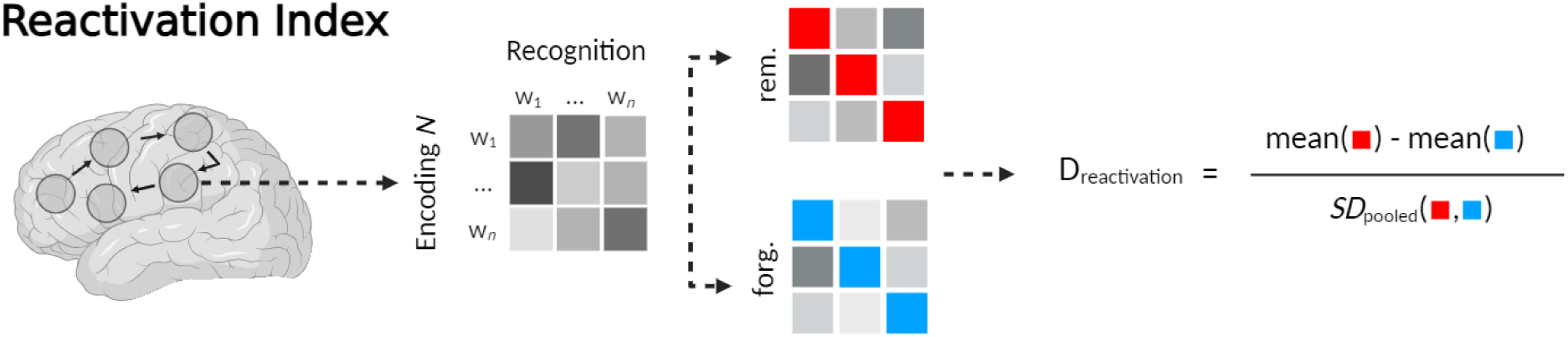
A reactivation index was computed at each searchlight centre. We first calculated pairwise correlations for each word (w_1 …_ *_n_*) between each encoding run (Encoding *N*) and the recognition run (Recognition). Cells (squares) in the grayscale matrices represent correlations between pairs of words; on-diagonal cells contain same-item correlations used in this analysis (off-diagonal cells are not relevant to this analysis, and were therefore not computed, but are shown in the illustration to facilitate conceptual understanding). Same-item correlations (on-diagonal cells) were sorted according to subsequent memory in the recognition phase: either subsequently remembered (rem.) or subsequently forgotten (forg.). A reactivation index (D_reactivation_) was computed as the standardised mean difference between correlations for correct items (red squares) versus correlations for incorrect items (blue squares).

#### 2.5.2. Transformation

We performed the transformation analysis in a manner that was as methodologically consistent as possible with the reactivation analysis. We broadly conceptualised transformation as the difference between encoding-recognition similarity and encoding-encoding (same item, between runs) similarity within each searchlight sphere. As such, we computed a *transformation index* as the difference (of encoding-recognition / encoding-encoding differences) between subsequently remembered and forgotten items—therefore, our transformation index was behaviourally relevant in the same way as our reactivation index.

Within each searchlight sphere, we first computed same-item correlations for encoding-encoding and encoding-recognition pairings, separately for subsequently remembered and forgotten items (as with the reactivation analysis, we considered correlations for both encoding runs relative to recognition). Following this, for each item, we subtracted each encoding-recognition correlation value from the encoding-encoding correlation value. The reasoning behind this step is that if patterns are transformed between encoding and recognition, encoding-recognition correlations should be smaller than encoding-encoding correlations, meaning that the product of the above subtraction should be a positive value. Moreover, the magnitude of this value should increase with larger differences between encoding-encoding and encoding-recognition (i.e., larger values correspond to greater degrees of transformation). This step generated two sets of values (one set per response type: subsequently remembered or forgotten), wherein every value reflected the difference in correlations between encoding-encoding and encoding-recognition for a single unique item. We computed a transformation index as the normalised mean difference (Cohen’s D) in correlation-difference values between subsequently remembered and subsequently forgotten items. This procedure is illustrated in Figure 3a. As before, we generated maps of transformation indices independently for each participant and condition. Hereon in, we will refer to this measure as the *within-subjects transformation index*.

**Figure 3.**
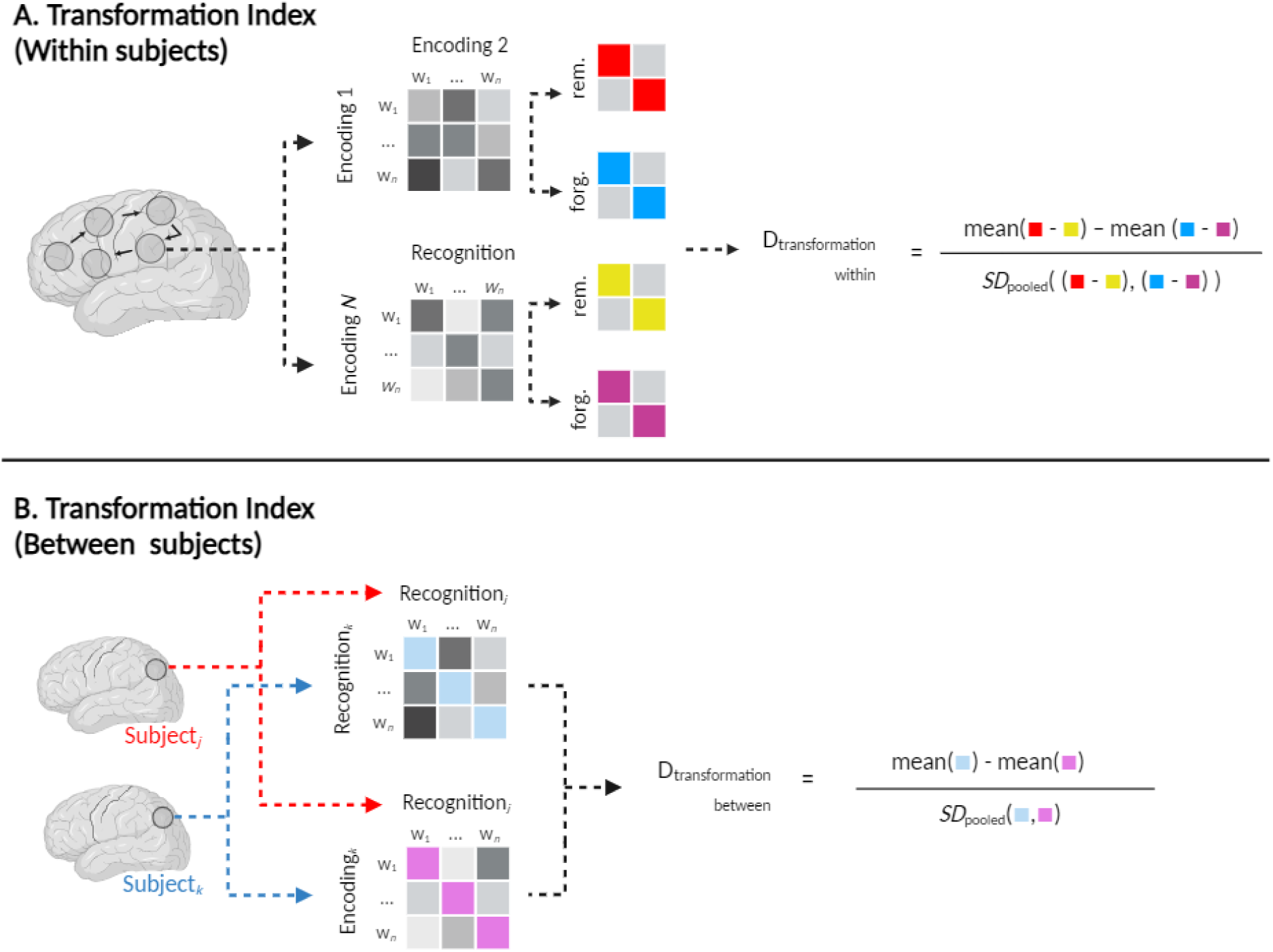
Within- and between-subjects transformation indices were calculated at each searchlight centre, for each individual subject. **A.** Within subjects: we calculated pairwise correlations for each word w_1 …_ *_n_*between the two encoding runs [*r*(Encoding 1, Encoding 2)], separately for subsequently remembered items (rem., red squares) and forgotten items (forg., blue squares). We also calculated pairwise correlations between each encoding run and the recognition run [*r*(Encoding *N*, Recognition)], again for remembered (gold squares) and forgotten (purple squares) items. Correlation difference values were computed as [*r*(Encoding 1, Encoding 2) - *r*(Encoding *N*, Recognition)], separately for remembered and forgotten items. A within-subjects transformation index (D_transformation within_) was then computed as the standardised mean difference between remembered correlation difference values and forgotten correlation difference values. **B.** Between subjects: activity patterns from each subject *k* were iteratively compared to every other subject *j*. This analysis only considered correctly remembered items that were shared by *k* and *j*. We computed same-item correlations for each word w_1 …_ *_n_* between encoding*_k_* (patterns were averaged across encoding runs) and retrieval*_j_*, and between retrieval*_k_*and retrieval*_j_*. A between-subjects transformation index (D*_t_*_ransformation between_) was computed as the standardised mean difference between *r*(recognition*_k_*, recognition*_j_*) and *r*(encoding*_k_*–recognition*_j_*).

To verify that our within-subjects index captured true transformation—that is, systematic changes in activation patterns, rather than mere weakening of encoding patterns or an increase in noise (Xue, 2022)—we constrained the results of our within-subjects analysis with those of a between-subjects analysis (similar to Chen et al., 2017). To be clear: results from the between-subjects analysis are not reported here: rather, they were simply used to mask results of the within-subjects analysis. The between-subjects analysis allowed us to identify areas in which recognition-recognition correlations (across participants) were stronger than encoding-recognition correlations (also across participants). If apparent transformation effects were driven by weakened / more noisy response patterns, stimuli should not be more decodable at recognition (Xue, 2022). Stated differently: if systematic transformation had occurred, patterns elicited (in different participants) by the same item during the recognition phase should be more highly correlated than patterns elicited between encoding and recognition.

We computed a *between-subjects transformation index* at each searchlight centre, for each participant. This index was based on correlations of activity patterns between pairs of participants (see below), therefore our calculations were necessarily constrained to items that were shared across pairs (which varied because words were randomly assigned to aloud or silent reading). For this reason, we only considered items that were subsequently remembered by both subjects in each pair; it was not feasible to compute a remembered > forgotten between-subjects index, as very few forgotten items were shared across participants.

First, for each participant, we iteratively computed same-item correlations between patterns at encoding^4^ or recognition from that (*k*th) participant and patterns at recognition from each *other* (*j*th) participant. This yielded between-participant values for *r*(encoding*_k_*, recognition*_j_*) and *r*(recognition*_k_*, recognition*_j_*). We then computed (for each participant) a between-subjects transformation index as Cohen’s D for the difference between *r*(recognition*_k_*, recognition*_j_*) values and *r*(encoding*_k_*, recognition*_j_*) values. This calculation is illustrated in Figure 3b.

### 2.6. Group-Level Analyses

#### 2.6.1. Behavioural Responses

Behavioural analyses were performed in the R environment (version 1.0.4; R Core Team, 2021). We analysed participants’ button-press responses from the recognition task. During this task, participants saw words that they had either read aloud or silently during the encoding phase, or had not previously seen (foil words). On each trial, participants indicated whether the presented word was “old” or “new”. Performance was measured as the percentage of recognition trials on which participants made an “old” response (as in Bailey et al., 2021); statistical analyses were performed on mean percentages of “old” responses in each condition from each participant. Statistical analysis between conditions was based on Bayes Factors (BFs), computed in the R environment using the BayesFactor package (Morey et al., 2022). A Bayes Factor BF_10_ is a ratio expressing the likelihood of the data under an alternative hypothesis (H_1_) relative to the null hypothesis (H_0_). Within this framework, increasing BF_10_ values > 1.0 correspond to increasing strength of evidence for H_1_ (Dienes, 2014; Lee & Wagenmakers, 2014). We tested the hypothesis H_1_ that the aloud condition elicited more “old” responses compared to the silent and foil conditions, with separate paired-samples two-tailed Bayes *t*-tests, using default JZS priors (Rouder et al., 2009); this yielded a single Bayes Factor for each between-condition contrast.

#### 2.6.2. Multivariate fMRI Responses

We performed group-level analyses on participants’ searchlight output based on feature (voxel)-wise Bayes Factors, in line with the application of Bayes Factors in previous multivariate neuroimaging studies (Kaiser et al., 2018; Grootswagers et al., 2019a, 2019b; Proklova et al., 2019; Teichmann et al., 2021; Moerel et al., 2022; Matheson et al., 2023). We performed voxel-wise Bayes *t*-tests in the MATLAB environment with functions from the bayesFactor package (Krekelberg, 2022), using default JZS priors (Rouder et al., 2009). We first tested for the presence of reactivation and transformation (both within- and between-subjects) in each condition alone. At each searchlight centre we performed a one-sample right-tailed Bayes *t*-test for the hypothesis H_1_ that reactivation/transformation indices were greater than zero. This procedure was repeated for each condition and measure, yielding six whole-brain statistical maps of BF_10_ values. Results from these within-condition analyses are not explicitly reported here; rather, they were used to constrain results from the between-condition analyses (described as follows). Next, we sought to test the hypothesis H_1_ that there was a difference in either reactivation or within-subject transformation between the aloud and silent conditions. At each searchlight centre, we performed a two-tailed Bayes *t*-test for a difference between conditions. This procedure was repeated independently for each measure, thus generating two statistical maps of BF_10_ values.

We thresholded all group-level statistical maps at BF_10_ ≥ 3.00, thereby constraining results to voxels expressing at least moderate evidence for H_1_ (consistent with conventional qualitative benchmarking of Bayes Factors; Lee and Wagenmakers, 2014). In the context of a whole-brain analysis, we do not feel that reporting weak evidence (BF_10_ < 3) would be particularly valuable, as it would distract from areas where there was relatively stronger evidence for H_1_. To determine directionality of between-condition effects concerning reactivation and within-subjects transformation, we calculated two average contrast maps for each of those measures. An aloud > silent map was computed by subtracting the average searchlight results (across all participants) in the silent condition from those of the aloud condition. A silent > aloud map was computed in the opposite direction. These average contrast maps were used to mask the thresholded BF_10_ maps, meaning that each measure was now associated with two BF_10_ maps: one reflecting voxels where group-average values were higher in the aloud condition compared to silent (aloud > silent), and one vice-versa (silent > aloud). We further constrained the between-condition BF_10_ maps by masking each with the one-sample *t*-test map for its respective minuend condition (e.g., the aloud > silent reactivation map was masked with the one-sample reactivation map for the aloud condition). This ensured that areas showing differences between conditions were not driven entirely by negative values in the subtrahend condition.

Finally, we masked each within-subjects transformation contrast (aloud > silent or silent > aloud) with the between-subjects map for its corresponding minuend condition. Our choice to perform masking with individual conditions (as opposed to contrasts between conditions) is based on the following reasoning. The between-subjects measure was intended to capture transformation as conceptualised in previous literature (i.e., not driven by noise or weakened responses) (Xue, 2022). It was *not* intended to capture behaviourally-relevant differences that might exist between aloud and silent reading. As such, this analysis simply served to verify whether areas implicated in the (behaviourally relevant) between-condition contrasts also exhibited “true” transformation in their respective minuend condition.

We generated interpretable tables of results for each contrast using FSL’s *cluster* and *atlasquery* functions, and custom bash scripting. Tables report spatial extent, mean and maximum BF_10_ values, and centre of gravity (COG) MNI coordinates for clusters of contiguous voxels (minimum spatial extent = 20 voxels). Mean and maximum BF_10_ are arguably both informative—local maxima are typically reported in fMRI studies, and in this context reflect the strongest available evidence for H_1_ in a given cluster. On the other hand, mean BF_10_ values are more representative of evidential strength when considering whole clusters. Anatomical labels for each cluster were identified from the Harvard-Oxford cortical and subcortical atlases (clusters whose COG fell outside anatomical structures defined by these atlases are not reported).

## 3. Results

### 3.1. Behavioural Responses

Condition-wise group means for the percentage of “old” responses on the recognition task are illustrated in Figure 4. The results of independent Bayes *t*-tests for differences between conditions revealed strong evidence for a higher proportion of “old” responses in the aloud condition relative to the silent (BF_10_ = 11.33) and foil (BF_10_ > 10,000) conditions.

**Figure 4.**
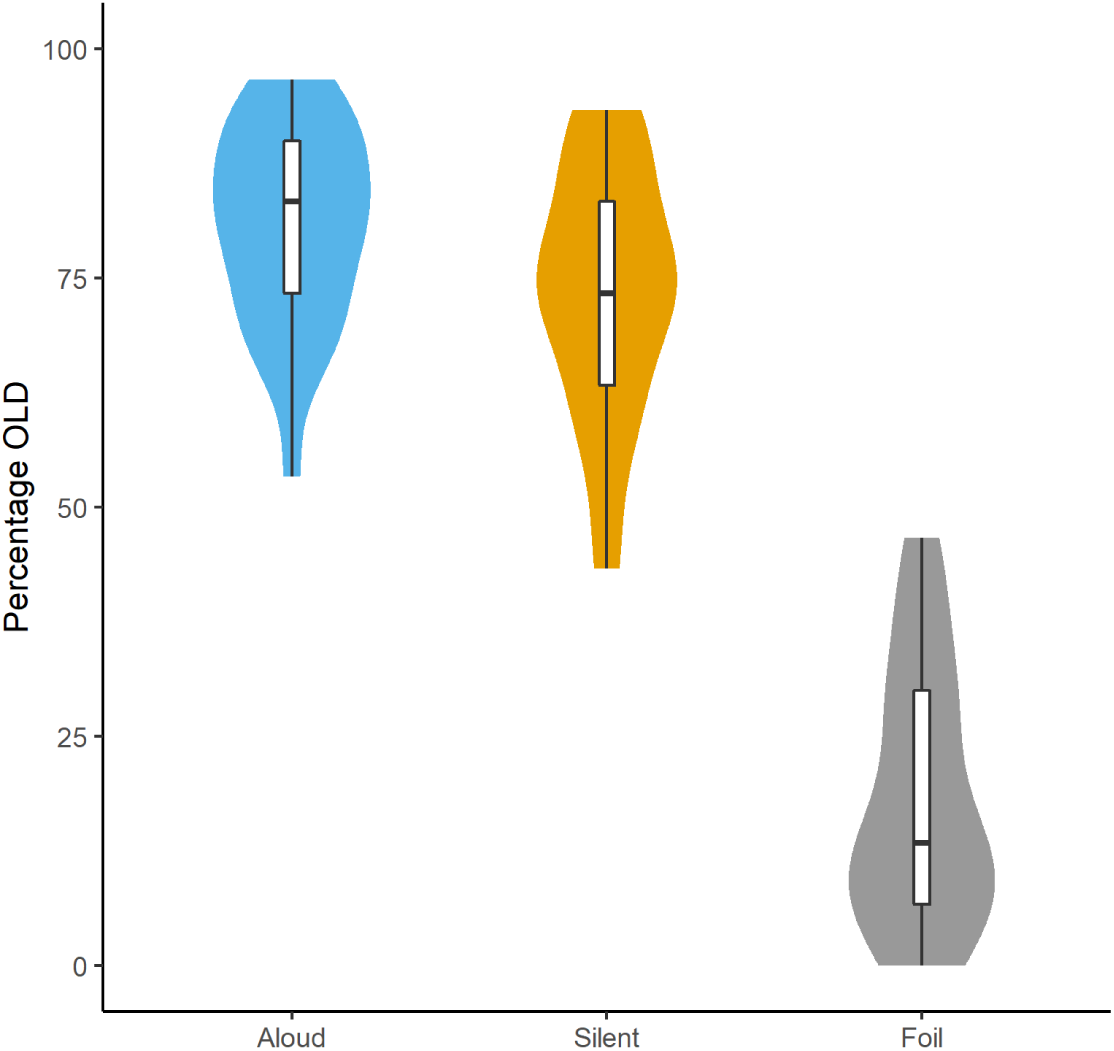
Percentages of OLD responses to words in each condition during the recognition task. Violin plots show the distributions of individual participant percentages in each condition. Box plots show condition-wise means across participants (middle bar), upper and lower quartiles (upper and lower box limits respectively) and ranges (whiskers).

### 3.2. Encoding-Recognition Reactivation

Results for the reactivation analysis are presented in Table 2 and Figure 5. The aloud > silent contrast revealed a single cluster with strong evidence (Mean, Maximum BF_10_ = 11.04, 38.40) situated in the posterior portion of the left insula.

**Figure 5.**
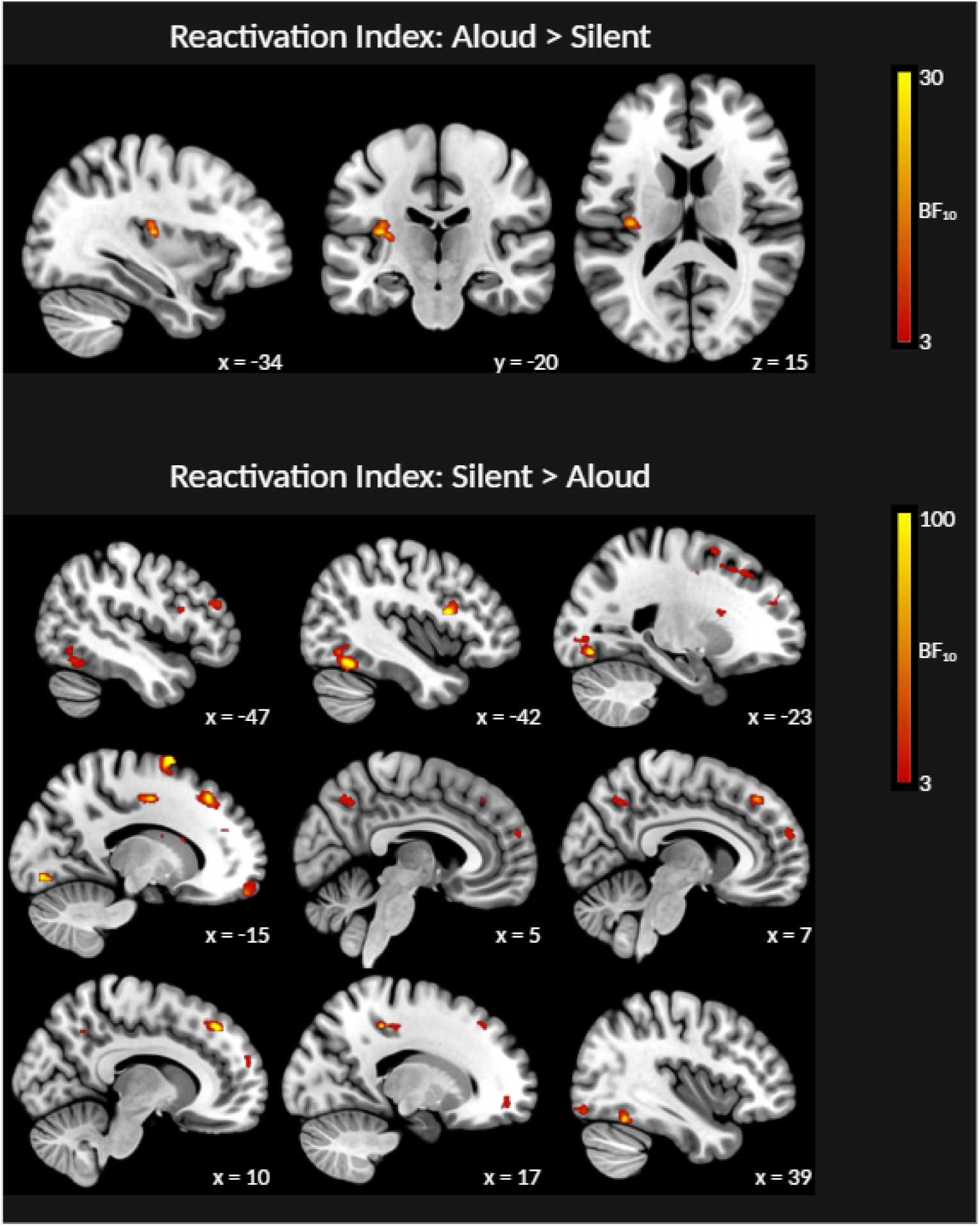
Cross-sectional slices show BF_10_ maps from the reactivation analyses. Highlighted areas show evidence of greater reactivation in the aloud reading condition relative to the silent reading condition (top panel) and vice-versa (bottom panel). Axial images are in neurological orientation (L-R).

**Table 2.**
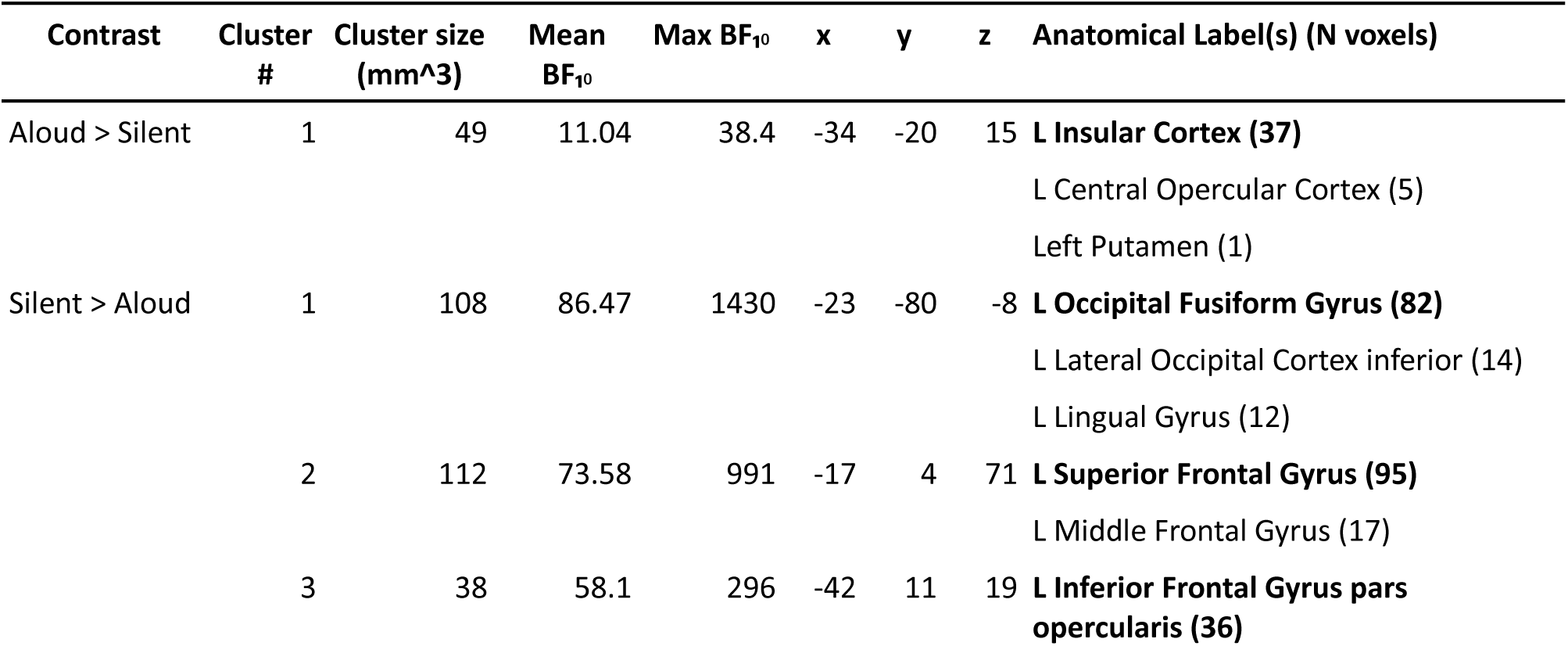
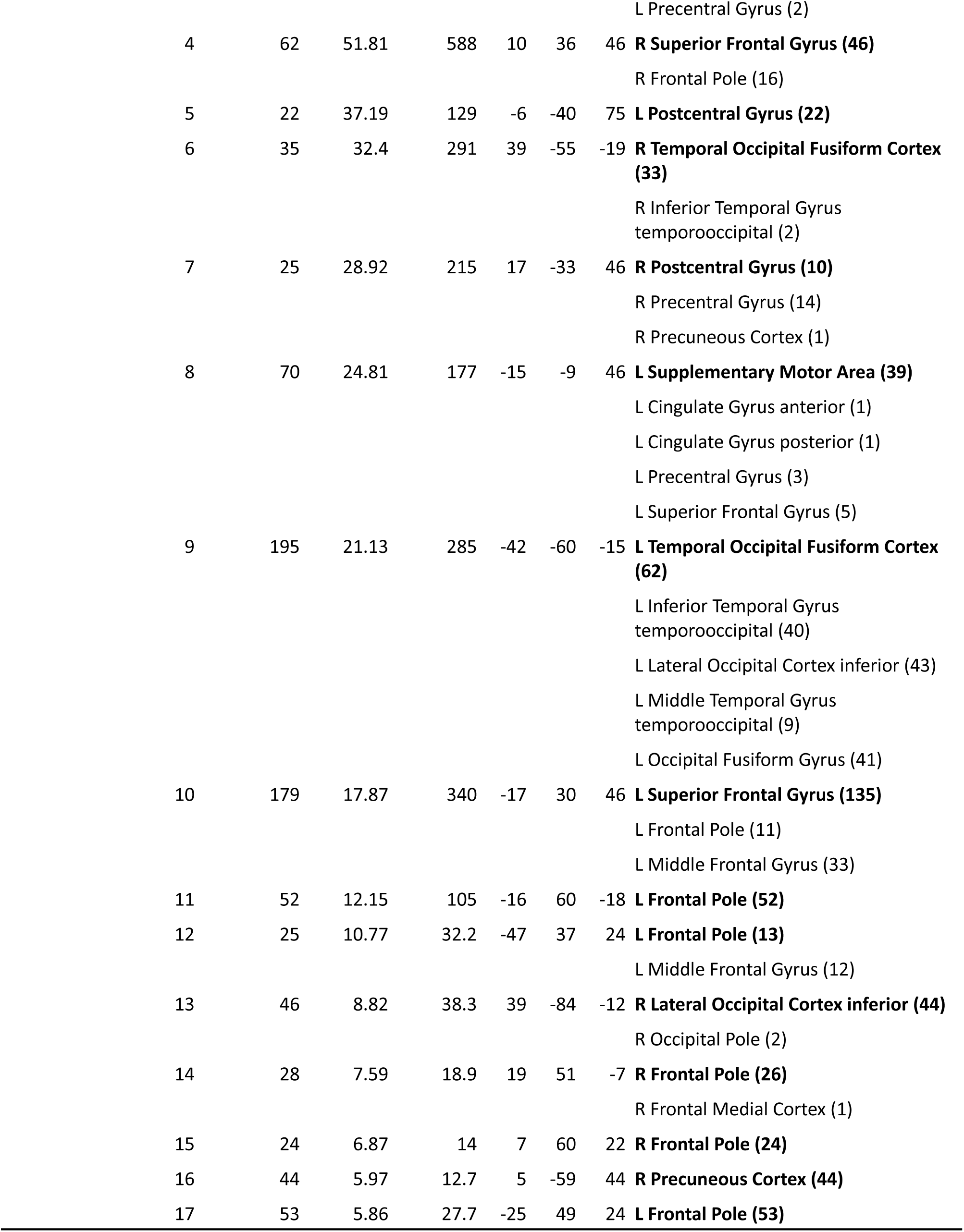
Clusters exhibiting evidence of between-condition differences in reactivation. Anatomical labels in boldface reflect the structure in which the centre of mass (COM) for that cluster was located.

| Contrast | Cluster # | Cluster size (mm<sup>3</sup>) | Mean $BF_{10}$ | Max $BF_{10}$ | x | y | z | Anatomical Label(s) (N voxels) |
| --- | --- | --- | --- | --- | --- | --- | --- | --- |
| Aloud > Silent | 1 | 49 | 11.04 | 38.4 | -34 | -20 | 15 | **L Insular Cortex (37)**<br>L Central Opercular Cortex (5)<br>Left Putamen (1) |
| Silent > Aloud | 1 | 108 | 86.47 | 1430 | -23 | -80 | -8 | **L Occipital Fusiform Gyrus (82)**<br>L Lateral Occipital Cortex inferior (14)<br>L Lingual Gyrus (12) |
|  | 2 | 112 | 73.58 | 991 | -17 | 4 | 71 | **L Superior Frontal Gyrus (95)**<br>L Middle Frontal Gyrus (17) |
|  | 3 | 38 | 58.1 | 296 | -42 | 11 | 19 | **L Inferior Frontal Gyrus pars opercularis (36)** |

|  |  |  |  |  |  |  |  |
| --- | --- | --- | --- | --- | --- | --- | --- |
|  |  |  |  |  |  |  | L Precentral Gyrus (2) |
| 4 | 62 | 51.81 | 588 | 10 | 36 | 46 | **R Superior Frontal Gyrus (46)** |
|  |  |  |  |  |  |  | R Frontal Pole (16) |
| 5 | 22 | 37.19 | 129 | -6 | -40 | 75 | **L Postcentral Gyrus (22)** |
| 6 | 35 | 32.4 | 291 | 39 | -55 | -19 | **R Temporal Occipital Fusiform Cortex (33)** |
|  |  |  |  |  |  |  | R Inferior Temporal Gyrus<br>temporooccipital (2) |
| 7 | 25 | 28.92 | 215 | 17 | -33 | 46 | **R Postcentral Gyrus (10)** |
|  |  |  |  |  |  |  | R Precentral Gyrus (14) |
|  |  |  |  |  |  |  | R Precuneous Cortex (1) |
| 8 | 70 | 24.81 | 177 | -15 | -9 | 46 | **L Supplementary Motor Area (39)** |
|  |  |  |  |  |  |  | L Cingulate Gyrus anterior (1) |
|  |  |  |  |  |  |  | L Cingulate Gyrus posterior (1) |
|  |  |  |  |  |  |  | L Precentral Gyrus (3) |
|  |  |  |  |  |  |  | L Superior Frontal Gyrus (5) |
| 9 | 195 | 21.13 | 285 | -42 | -60 | -15 | **L Temporal Occipital Fusiform Cortex (62)** |
|  |  |  |  |  |  |  | L Inferior Temporal Gyrus<br>temporooccipital (40) |
|  |  |  |  |  |  |  | L Lateral Occipital Cortex inferior (43) |
|  |  |  |  |  |  |  | L Middle Temporal Gyrus<br>temporooccipital (9) |
|  |  |  |  |  |  |  | L Occipital Fusiform Gyrus (41) |
| 10 | 179 | 17.87 | 340 | -17 | 30 | 46 | **L Superior Frontal Gyrus (135)** |
|  |  |  |  |  |  |  | L Frontal Pole (11) |
|  |  |  |  |  |  |  | L Middle Frontal Gyrus (33) |
| 11 | 52 | 12.15 | 105 | -16 | 60 | -18 | **L Frontal Pole (52)** |
| 12 | 25 | 10.77 | 32.2 | -47 | 37 | 24 | **L Frontal Pole (13)** |
|  |  |  |  |  |  |  | L Middle Frontal Gyrus (12) |
| 13 | 46 | 8.82 | 38.3 | 39 | -84 | -12 | **R Lateral Occipital Cortex inferior (44)** |
|  |  |  |  |  |  |  | R Occipital Pole (2) |
| 14 | 28 | 7.59 | 18.9 | 19 | 51 | -7 | **R Frontal Pole (26)** |
|  |  |  |  |  |  |  | R Frontal Medial Cortex (1) |
| 15 | 24 | 6.87 | 14 | 7 | 60 | 22 | **R Frontal Pole (24)** |
| 16 | 44 | 5.97 | 12.7 | 5 | -59 | 44 | **R Precuneous Cortex (44)** |
| 17 | 53 | 5.86 | 27.7 | -25 | 49 | 24 | **L Frontal Pole (53)** |

The silent > aloud contrast revealed more extensive reactivation, with clusters throughout frontal, parietal, and ventral temperooccipital cortices. In the frontal lobe, clusters with strong evidence (Mean BF_10_ ≥ 10) for this contrast were present in the superior and middle frontal gyri bilaterally, as well as the left pars opercularis and supplementary motor area. Frontal clusters with relatively weaker evidence (that is, Mean BFs closer to or < 10) were present in the frontal poles bilaterally. In the parietal lobe, clusters with strong evidence were present in the postcentral gyrus bilaterally; one additional parietal cluster with moderate evidence was situated in the right precuneus. In ventral temperooccipital cortex, clusters with strong evidence were present bilaterally, in the temperooccipital and (more posterior) occipital portions of the fusiform gyrus. Notably, the cluster with the strongest evidence for this contrast was situated in left occipital fusiform cortex (Mean, Maximum BF_10_ = 86.47, 1430).

### 3.3. Encoding-Recognition Transformation

Results for the transformation analysis are presented in Table 3 and Figure 6. The aloud > silent contrast revealed one cluster with strong evidence (Mean, Maximum BF_10_ = 991.56, 28000), situated in the posterior portion of the left precuneus. Another cluster with relatively weaker evidence (Mean, Maximum BF_10_ = 6.46, 11.70) was situated in the posterior portion of left ventral temporal cortex. The silent > aloud contrast did not reveal any clusters.

**Figure 6.**
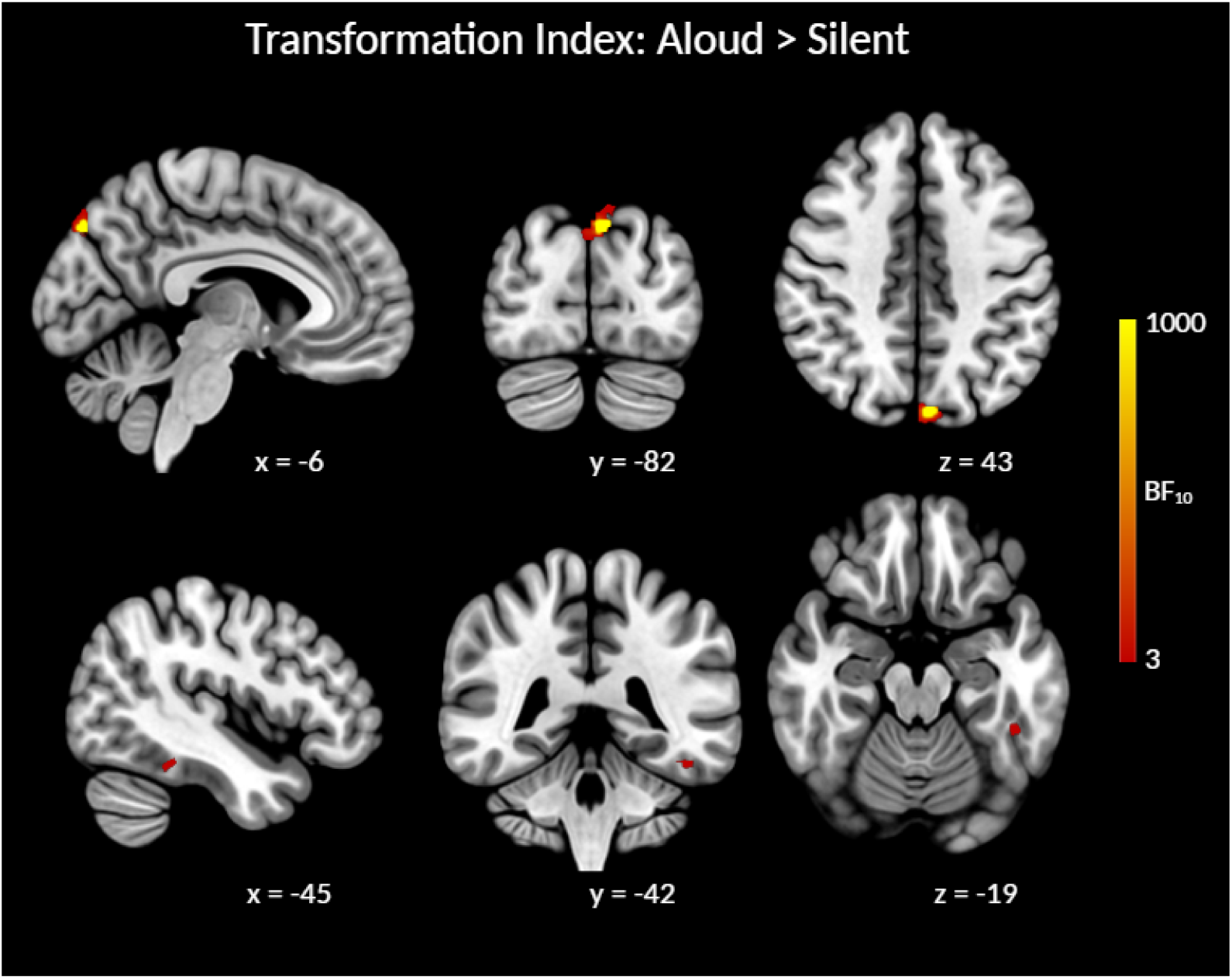
Cross-sectional slices show the BF_10_ map from the aloud > silent contrast of within-subjects transformation indices, masked with the map from the aloud > 0 contrast of between-subjects transformation indices. Highlighted areas show evidence of greater transformation in the aloud reading condition relative to the silent reading condition. Axial images are in neurological orientation (L-R).

**Table 3.** Clusters exhibiting evidence of between-condition differences in transformation. Anatomical labels in boldface reflect the structure in which the centre of mass (COM) for that cluster was located.

| Contrast | Cluster # | Cluster size (mm<sup>3</sup>) | Mean $BF_{10}$ | Max $BF_{10}$ | x | y | z | Anatomical Label(s) (N voxels) |
| --- | --- | --- | --- | --- | --- | --- | --- | --- |
| Aloud > Silent | 1 | 59 | 991.56 | 28000 | -6 | -82 | 43 | **L Precuneous Cortex (17)** |
|  |  |  |  |  |  |  |  | L Cuneal Cortex (7) |
|  |  |  |  |  |  |  |  | L Lateral Occipital Cortex superior (33) |
|  |  |  |  |  |  |  |  | R Cuneal Cortex (2) |
|  | 2 | 26 | 6.46 | 11.7 | -45 | -42 | -19 | **L Inferior Temporal Gyrus posterior (13)** |
|  |  |  |  |  |  |  |  | L Inferior Temporal Gyrus temporooccipital (1) |
|  |  |  |  |  |  |  |  | L Temporal Fusiform Cortex posterior (11) |
|  |  |  |  |  |  |  |  | L Temporal Occipital Fusiform Cortex (1) |

## 4. Discussion

This study builds upon prior work exploring neural correlates of the production effect (Hassall et al., 2016; Bailey et al., 2021; Zhang et al., 2023), and is the first such study to offer insights about its underlying spatial dynamics from a multivariate perspective. Our behavioural analyses showed that we replicated the standard production effect: participants remembered a larger proportion of words from the aloud condition relative to the silent condition. With respect to fMRI analyses, we investigated relative contributions of neural pattern reactivation and transformation during recognition of words from the aloud and silent reading conditions. Our reactivation and transformation measures quantified differences between subsequently remembered and forgotten items, therefore they inherently capture processes underlying recognition success. We compared reactivation and transformation between the two reading conditions, thus revealing areas in which these processes were differentially involved in remembering.

We predicted that we would observe relatively more reactivation and/or transformation for aloud words relative to silent words. While this was true for transformation—which was exclusively detected by the aloud > silent contrast—we in fact observed relatively more widespread reactivation in the silent > aloud contrast (evidenced by a larger number or clusters with generally stronger evidence) compared with the aloud > silent contrast. These results indicate that remembering words that were read aloud versus silently seems to depend, to some extent, on dissociable neural mechanisms. This is in contrast to current theoretical accounts of the production effect which are (necessarily) one sided; these accounts generally frame encoding and subsequent retrieval as being stronger or more elaborate for aloud words (e.g., Jamieson et al., 2016; MacLeod and Bodner, 2017; Fawcett et al., 2022), with little consideration of processes that might be enhanced for silent words. Stated differently, aloud reading is generally considered to entail all the same processes as silent reading, *plus* processes elicited by speech production. Our results provide a rather different perspective: that word recognition may entail different mechanisms, depending on the preceding reading conditions, rather than greater or lesser degrees of the same mechanism(s). Below we elaborate on the functional significance of reactivation and/or transformation effects identified by each contrast.

### Aloud > Silent Contrasts

The aloud > silent contrasts are of particular theoretical value, because they reveal apparent neural correlates of the production effect (defined as a behavioural contrast of aloud > silent). Our results here identified contributions of both reactivation and transformation, primarily in posterior portions of the left insula and precuneus respectively. The left insula has been linked to speech and language processes broadly (see Oh et al., 2014 for meta-analysis), while Woolnough et al. (2019) suggested that its posterior portion might mediate integration of somatosensory and auditory information during speech production. As such, reactivation in this area likely reflects reinstatement of speech-elicited processes, consistent with a distinctiveness-based account of the production effect (MacLeod et al., 2010; Jamieson et al., 2016).

Understanding the observed transformation effect in the precuneus is less straightforward, and requires careful consideration of the precuneus’ known functions. This area has been associated with a number of processes, notably including episodic memory retrieval and metacognition. With respect to the former, fMRI work has implicated posterior precuneus in recollection (i.e., the subjective experience of remembering a specific event, as opposed to the more general sensation of familiarity) (Henson et al., 1999; Fandakova et al., 2021), memory for contextual details (e.g., source memory judgements; Lundstrom et al., 2003, 2005), and vivid reminiscence (Richter et al., 2016). The precuneus has also been linked to memory-relevant metacognition—that is, evaluation of one’s internal cognitive state as it pertains to mnemonic decisions. One study reported that grey matter concentration in posterior precuneus was correlated with participants’ ability to appropriately evaluate their own performance on a memory task (selecting a previously studied stimulus from two alternatives; McCurdy et al., 2013), while self-evaluation during a similar task also elicited activation in the precuneus (albeit in a more inferior portion) relative to a perceptual judgement task (Morales et al., 2018). Moreover, a TMS study reported that selective disruption of the precuneus perturbed participants’ ability to appropriately evaluate their own memory (Ye et al., 2018). In a slightly different vein, the paracingulate network—which is contained within the precuneus, and importantly includes its posterior portion (Dadario and Sughrue, 2023)—is uniquely suited to incorporate speech information as a component of both recollection and metacognition. This network receives input from multiple areas, notably including the pre/postcentral gyri, insular, and supplementary motor area; all of which are involved in speech production (e.g., Bailey et al., 2021). Dadario and Sughrue (2023) suggest that this network acts as a hub for integrating external sensory information (via functional connections to the aforementioned areas) with internal knowledge (e.g., introspective information and knowledge of task goals) for the purpose of guiding goal-directed behaviour. Therefore, we reason that the precuneus may have the capacity to incorporate encoded sensorimotor information (that was elicited during speech production) during memory-related metacognition, particularly in the context of a specific cognitive goal (e.g., deciding whether a presented word is old or new).

Based on the above information, we suggest that transformation in the precuneus may reflect access to, and subsequent metacognitive evaluation of encoded speech-related information, for the purpose of making old-new judgments during the recognition task. This encoded (and subsequently evaluated) information may be received as input from the insula, which is functionally connected to the paracingulate network (Dadario and Sughrue, 2023) and which exhibited reactivation effects in our analysis. Our proposal is consistent with the notion that participants’ recognition judgements on aloud words are guided by an evaluative heuristic (MacLeod and Bodner, 2017); in this case, however, remembering is not entirely guided by faithful reinstatement (“replaying”) of encoded experiences. As discussed in the introduction, evaluating encoded information, as opposed to mentally replaying it, may require reorganisation of that information (consistent with prior multivariate studies showing goal-dependent changes in neural decodability; Nastase et al., 2017; Favila et al., 2018; Wang et al., 2018).

An alternative interpretation (though one which is not mutually exclusive with a heuristic account) is that transformation in the precuneus may reflect some component(s) of the retrieval process described by Wakeham-Lewis et al. (2022). By this account, when presented with a word (for which they must make an old/new decision), participants may simulate reading the presented word aloud and then compare the simulated experience with available encoded information associated with that word. It seems feasible that the precuneus might play a role here: in addition to its roles in recollection and metacognition, the precuneus is reportedly activated by a number of tasks involving mental simulation from a first-person perspective, such as mental imagery and imagined navigation (see Cavanna and Trimble, 2006 for review). More recently, Tanaka and Kirino (2021) report that during imagined singing, the precuneus exhibited heightened functional connectivity with perisylvian areas associated with speech production (inferior frontal and temporal gyri), as well as the middle and (medial) superior frontal gyri, which have both been linked to planning and cognitive control of speech output (Bourguignon, 2014; Hertrich et al., 2016, 2021). Takana and Kirino (2021) suggest that their results may reflect integration of language-related information during imagined singing. With respect to our study, because singing bears perceptual/experiential similarities to reading aloud (vocalisation and sensory feedback), it seems feasible that the precuneus would play a similar role in imagined aloud reading. Having said this, it remains unclear whether our observed transformation effect reflects simulation *per-se*, or the process of comparing the simulated experience with the encoded experience (Wakeham-Lewis et al., 2022). As discussed above, the metacognitive functional properties of the precuneus seem compatible with evaluation of encoded sensorimotor information; in this case, comparing mental simulation to past experiences. As before, such evaluation may be possible owing to the precuneus’ functional connections with language-related areas involved in the initial productive experience. Considering that (by Wakeham-Lewis and colleagues’ account) simulation should take place on *all* recognition trials (including both aloud and silent words), our aloud > silent transformation effect may reflect successful matching between the simulated speech and encoded information on (correctly recognised) aloud trials. Unfortunately, it is difficult to separate mental simulation from comparison (to encoded information) based on the current data; we feel that further research is required to properly address this issue.

### Silent > Aloud Contrasts

The silent > aloud reactivation contrast revealed a number of clusters; notably, clusters with the strongest evidence were situated in posterior fusiform and prefrontal cortices. Prior work has indicated that posterior fusiform cortex is sensitive to low-level orthographic properties of printed words (e.g., consonants versus false fonts) (Vinckier et al., 2007; Thesen et al., 2012); its role in our study likely reflects reinstatement of these visual features during successful recognition. This finding is consistent with previous multivariate work reporting reactivation in early visual areas during encoding and retrieval of images (e.g., Bosch et al., 2014; Bone and Buchsbaum, 2021). We note that an adjacent cluster in the left temperooccipital portion of the fusiform gyrus (Cluster #9 in Table 2) approximately corresponds to the visual word-form area (VWFA)^5^, which has been the focus of much work surrounding the neural correlates of reading. The VWFA is generally considered central to comprehension of visual word-forms (e.g., Petersen et al., 1990; Cohen et al., 2002; McCandliss et al., 2003; Brem et al., 2010; Carreiras et al., 2014; Turkeltaub et al., 2014)—that is, perceptually-invariant mental representations of printed words (Warrington and Shallice, 1980). Our finding that this area (in addition to posterior fusiform gyrus) demonstrated reactivation for silent words may signal greater dependence (relative to aloud words) on orthographic information during recognition, likely because the initial encoding experience was entirely visual.

With respect to prefrontal areas, prior work has implicated both the pars opercularis and supplementary motor area in processes necessary for speech planning and production. In the context of single word reading, the pars opercularis has traditionally been associated with grapheme-to-phoneme mapping (Fiez et al., 1999; Mechelli et al., 2005), while supplementary motor area is thought to play a role in cognitive control of speech-related motor processes such as initiation and timing (Hertrich et al., 2016). As for the superior frontal gyrus, this area has been linked to domain-general cognitive control, particularly cognitive flexibility (i.e., task- or response rule-switching) and response inhibition (e.g., Konishi et al., 2003; Cutini et al., 2008; see Niendam et al., 2012 for a meta-analysis of these processes). Given the appearance of reactivation in multiple areas that are either involved in speech planning or cognitive control, we suggest that prefrontal reactivation might reflect reinstatement of (item-specific) inhibitory ‘codes’ related to suppression of vocal responses during silent reading. These codes may facilitate a recognition heuristic similar to that described by Macleod and Bodner (2017), but in this case signalling that words were *not* produced (*I remember stopping myself from reading this word aloud, therefore I must have seen it before*).

## 5. Conclusion

This study was the first to investigate neural pattern reactivation and transformation during recognition in the context of the production effect. Our results broadly support a distinctiveness-based account whereby recognising aloud words depends on retrieval of item-specific speech information. That being said, this conclusion rather depends on how one conceptualises retrieval. Speech information may be reinstated in its original encoded format, consistent with the idea of mentally “replaying” sensorimotor experiences (MacLeod and Bodner, 2017). On the other hand, the encoded information may be manipulated or reorganised, perhaps reflecting metacognitive evaluation of the recollected (or simulated) experience. Recognising silent words seems relatively more dependent on reinstatement. This may be because, unlike aloud words, silent words were not encoded alongside the unique sensorimotor experiences associated with articulation; therefore, participants must rely more heavily on visual-orthographic information.

## Data Availability

Code for all analyses reported in this manuscript is available on GitHub: https://github.com/lbailey25/Production_Effect_MVPA; additional materials that are necessary for analyses are stored in an Open Science Framework (OSF) repository: https://osf.io/czb26/?view_only=86a66caf1d71484d8ef0293cfa2371df. Twelve participants consented to their anonymized data being made publicly available. Raw data from those participants are available on the OSF repository. Note that the data reported in this manuscript are from the “PE” experiment described in both repositories.

## CRediT authorship statement

**Lyam M. Bailey:** Conceptualization, Methodology, Software, Formal analysis, Investigation, Resources, Data curation, Writing - original draft, Visualization, Project administration, Funding acquisition. **Heath E. Matheson:** Conceptualization, Methodology, Software, Writing - review & editing. **Jonathan M. Fawcett:** Conceptualization, Writing - review & editing. **Glen E. Bodner:** Conceptualization, Funding acquisition. **Aaron J. Newman:** Conceptualization, Methodology, Supervision, Project administration, Funding acquisition.

## Footnotes

1 This method parcellates a large area of cortex (in this case, the whole brain) into a series of smaller searchlight areas, and examinnes activation patterns within each area. As such, it allows one to effectively “scan” the entire brain, as opposed to relying on a set of discrete ROIs.

2 The only exception was the word *trousers* from Bailey et al. (2021); during piloting for the current study, multiple participants reported that this word “stood out” as unusual. Therefore, we replaced this word with *pants* in the current study.

3 We added 4 axial slices (total = 38) to the protocol for one participant, in order to accommodate their entire cerebral cortex.

4 To reduce computational load, for each participant we first averaged together activity patterns from the two encoding runs for each item. As such, this analysis did not include separate correlations for each encoding run, as did the reactivation and within-subjects transformation analyses.

5 Multiple studies (e.g., Vogel et al., 2012; Martin et al., 2015; Chen et al., 2019) consider the putative VWFA to be centred on the following coordinates: x = −45, y = −57, z = −12. The COG for the cluster we identified in temperooccipital fusiform cortex was x = −42, y = −60, z = −15; within 3 voxels of putative VWFA in any direction

